# Error-correcting dynamics in visual working memory

**DOI:** 10.1101/319103

**Authors:** Matthew F. Panichello, Brian DePasquale, Jonathan W. Pillow, Timothy J. Buschman

## Abstract

Working memory is critical to cognition, decoupling behavior from the immediate world. Yet, it is imperfect; internal noise introduces errors into memory representations (1, 2). Such errors accumulate over time (3–5) and increase with the number of items simultaneously held in working memory (6–10). Here, we show that error-correcting attractor dynamics mitigate the impact of noise on working memory. These dynamics pull memories towards a few stable representations in mnemonic space, inducing a bias in memory representations but reducing the effect of noise. Model-based and model-free analyses show attractor dynamics account for the frequency, bias, and precision of working memory reports in both humans and monkeys. Furthermore, attractor dynamics were optimized to the context; they adapted to the statistics of the environment, such that memories drifted towards contextually-predicted values. Our results suggest attractor dynamics mediate errors in working memory by counteracting noise and integrating contextual information into memories.

## Main Text

To understand the dynamics of errors in working memory, we examined the behavior of humans (N=90) and monkeys (N=2) performing a delayed continuous report working memory task (11; Fig. 1a). Subjects were instructed to remember the color of 1 to 3 simultaneously-presented stimuli located at different positions on the display (humans saw 1 or 3 items; monkeys saw 1 or 2). After a variable memory delay, subjects reported the remembered color at a cued target location using a continuous scale. Stimulus colors were drawn uniformly from an isoluminant circular color space. We quantified error as the angular deviation between the target color and the subject’s report. As expected (3–10), the average absolute error increased as a function of delay and working memory load (Fig. 1b; humans (H): load, *F* (1, 89) = 147.23, *p* < 0.001; delay, *F* (1, 89) = 85.44, *p* < 0.001; load x delay, *F* (1, 89) = 13.92, *p* < 0.001; monkey W (W): load, *p* < 0.001; delay, *p* = 0.006; load x delay, *p* = 0.495; monkey E (E): load, *p* < 0.001; delay, *p* = 0.009; load x delay, *p* = 0.303).

**Fig. 1:**
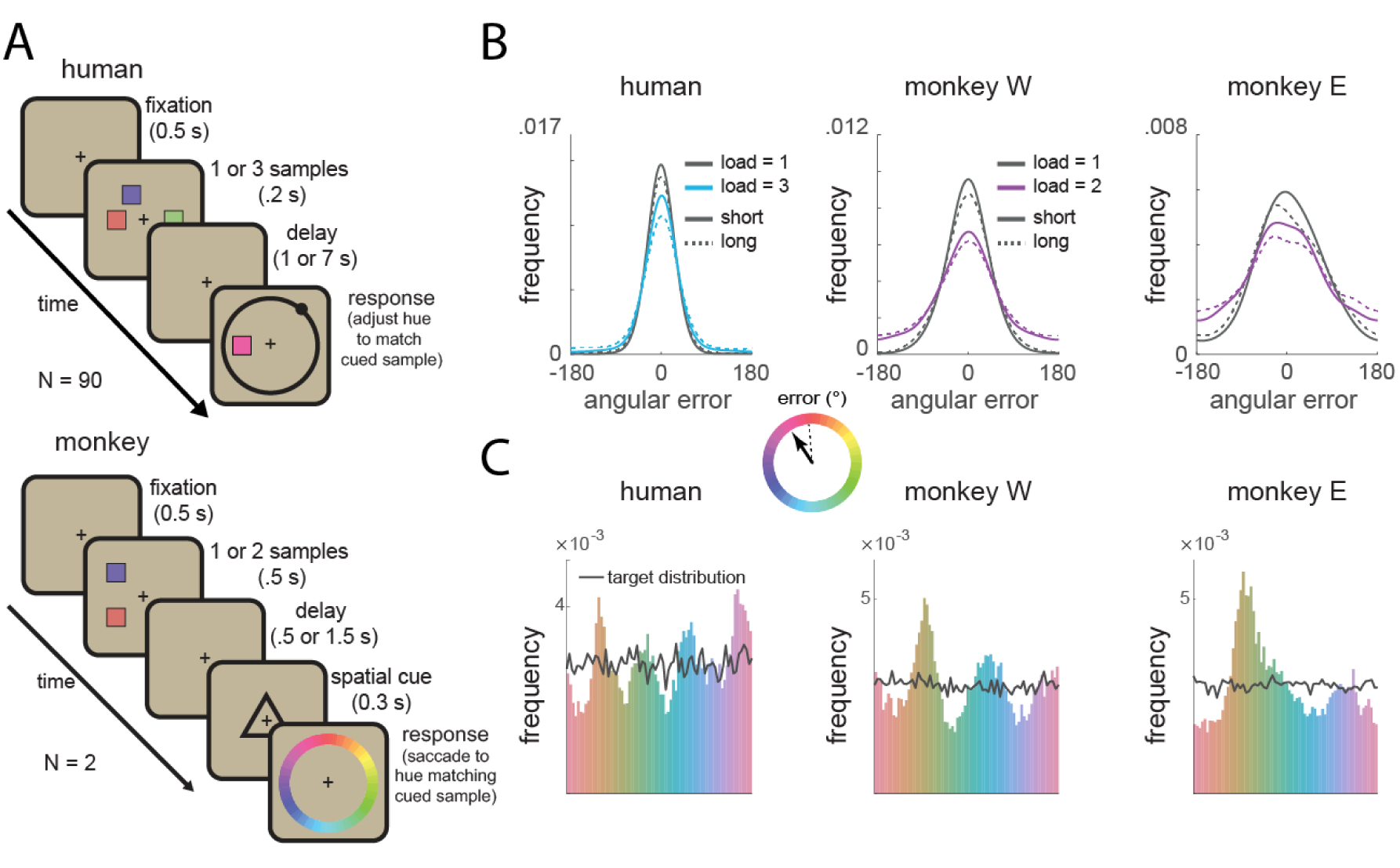
Experiment 1 task and behavioral performance. (A) Top: humans (N=90) performed a color delayed-estimation task in which they reported the color of a spatially-cued sample after a variable delay. Humans made their report by adjusting the hue of the response probe by rotating a response wheel (black circle) using a mouse. We rotated the mapping between wheel angle and color on each trial to avoid spatial encoding of color memories. Bottom: monkeys (N=2) performed a similar task. A symbolic cue indicated which sample to report (top or bottom). Monkeys reported a specific color value using an eye movement to a color wheel that was rotated on each trial. (B) Distribution of angular error for humans (top) and monkeys (bottom). Error increased with load and delay time. Inset: Error is calculated as the angular deviation between the color of the cued sample and the reported color in color space. (C) Non-uniform distribution of reported colors for humans (top) and monkeys (bottom). Black line shows the frequency of target colors.

Despite the uniform distribution of target colors, the responses of both human and monkey subjects were significantly non-uniform (12–15; humans and monkeys *p* < 0.001 against uniformity, Hodges-Ajne test; *p* < 0.001 against target distribution, permuted Kuiper’s test). This was reflected in a significant decrease in the entropy of the response distribution relative to the target distribution (H: 2.54 vs. 2.61 bits, *t*(89) = 13.90, *p* < 0.001, t-test; W: 2.61 vs. 2.65 bits, *p* < 0.001, bootstrap; E: 2.58 vs. 2.65 bits, *p* < 0.001, bootstrap). Responses clustered around specific colors, seen as peaks in the response histogram (Fig. 1c). Clustering increased with delay time (*F* (1, 89) = 9.56, *p* = 0.003) and with memory load in humans (Figs. S1 and S2; *F* (1, 89) = 5.45, *p* = 0.022). These results suggest clustering is the result of a load-dependent dynamic process.

Motivated by these results, we tested the hypothesis that attractor dynamics underlie the evolution of working memory representations. Attractors can be conceptualized as local minima in an ‘energy landscape’ over mnemonic (color) space, such that memories drift towards nearby attractors over time (Fig. 2a). This would provide a mechanistic explanation for the observed clustering of memory reports.

**Fig. 2:**
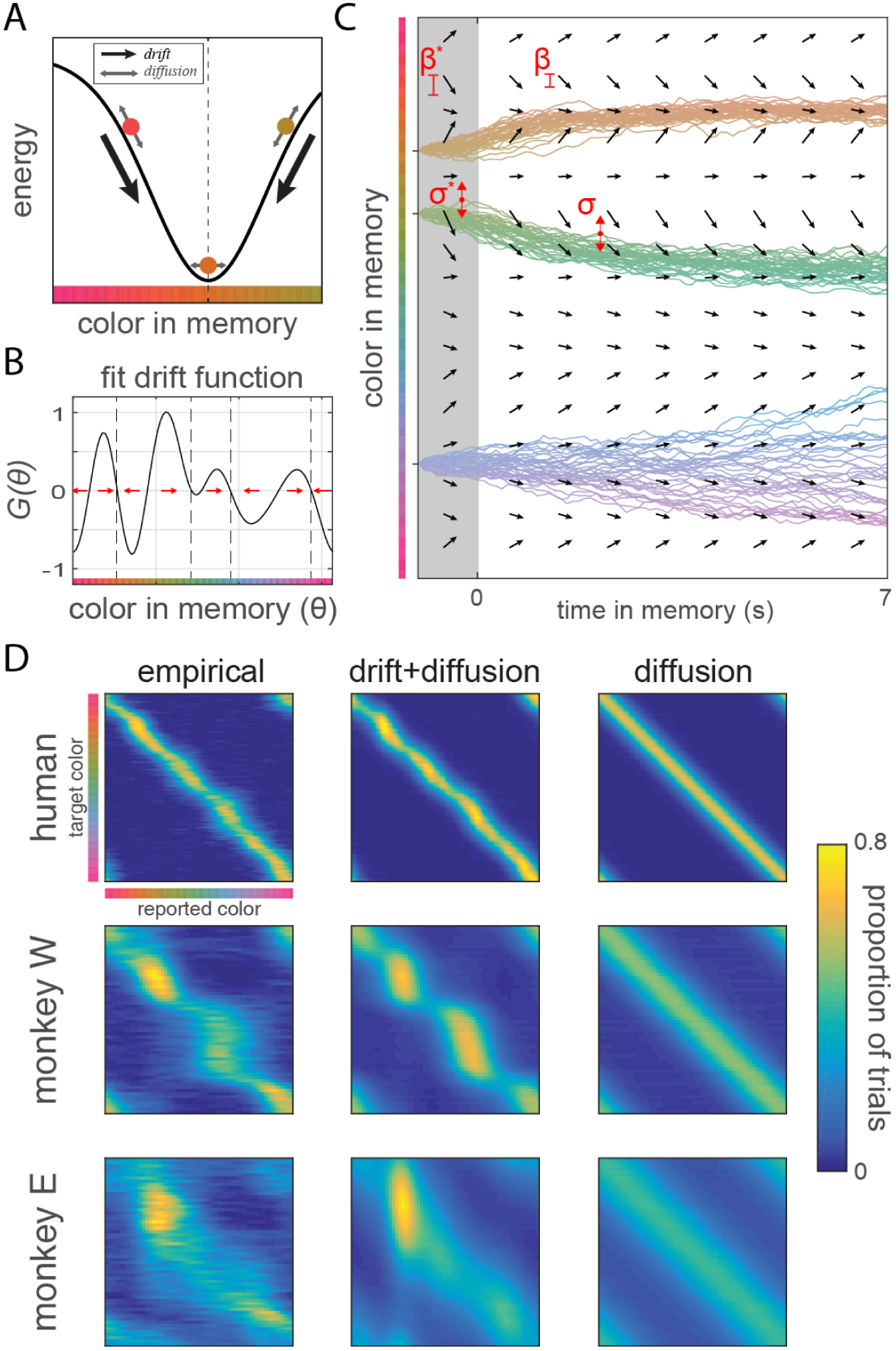
Dynamical model structure and performance. (A) Illustration of the influence of attractors on color memory. Attractors (dashed line) cause memories to drift over time (black arrow), introducing bias in reports. Noise causes memories to randomly diffuse (grey arrows). (B) The drift function *G*(*θ*) describes how a memory will change based on its current state. Red arrows show the direction of drift; attractors have converging drift (dashed lines). We estimated *G*(*θ*) for each subject using a linear combination of von mises derivatives. (C) The simulated evolution of three color memories during a hypothetical trial based on an example subject’s fit dynamical model. Memory evolves over time according to the drift function (vector field) and random noise. Each line indicates the temporal evolution of a remembered color under a different realization of the noise process. Terms described in main text. (D) Probability of responses (x-axis) for each target color (y-axis). The distribution of observed responses is better fit by the drift+diffusion model (middle) than by a model with only diffusion (right).

To test for the existence of attractors, we developed a dynamical systems model to characterize the dynamics governing working memory representations. The model describes memory error as due to a combination of ‘diffusion’ from noise in the neural representation (1, 16, 17) and ‘drift’ towards attractor states. Diffusion was quantified as a random walk from the current location in mnemonic space with no bias (*μ* = 0) and a variance 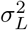 that depended on the number of colors presented (*L* = ‘memory load’). Attractor dynamics were estimated by fitting a drift function *G*(*θ*) for each subject that describes how a remembered color *θ* will drift as a function of its current value (Fig. 2b). Positive drift values reflect a clockwise drift (to the right in Fig. 2b) while negative values reflect a counterclockwise drift (to the left). Thus, attractors are points in mnemonic space that 1) are stable, such that they have no drift, and 2) pull nearby memories towards themselves, reflected in a negative slope in the local drift function (Fig. 2b, dashed lines). As with diffusion, the strength of the drift *β_L_* was load-dependent.

Together, drift and diffusion define the dynamic evolution of memories during the memory delay (Fig. 2c, *dθ* = *β_L_G*(*θ*)*dt* + *σ_L_* 𝒩(0, *dt*)). To capture errors during encoding (2, 18), inputs were first passed through an ‘encoding stage’ with the same drift and diffusion process, but with independent weighting parameters (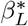 and 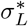). Finally, the model captured errors due to forgetting of memories (6) and responses to non-targets (7, see Methods for details). Model parameters were estimated by maximizing the likelihood of the observed memory reports for each subject. Critically, the model did not assume attractor dynamics (Fig. S3); when *β* and *β*^*^ are zero then memories are only influenced by diffusion, forgetting, and responses to non-targets, as in previous models.

Consistent with the hypothesis that attractor dynamics influence working memory, models that include drift fit significantly better than models without drift in both humans and monkeys (Fig. 2d and Fig. S4 and section; H: relative likelihood of full model = 1.00 (AIC) and 1.00 (BIC); W: 1.00 and 0.8; E: 1.00 and 1.00). The dynamical model accounted for 93.2% of the explainable variance in responses in humans and 87.7% and 83.7% in monkey W and E, respectively. This was significantly greater than a model that included diffusion alone (H: +5.3%, *p* < 0.001, bootstrap; W: +9.4%, *p* = 0.037, bootstrap; E: +19.6%, *p* = 0.006, bootstrap).

Attractor dynamics explain the frequency, bias, and precision of memory reports. First, attractors explain the observed clustering of memory reports. Indeed, the location of attractors identified using the drift function coincided with peaks in the response histogram. Fig. 3a shows the frequency of responses around attractors was significantly greater than that expected by chance (H: *t*(89) = 18.53, *p* < 0.001; W: *p* < 0.001, bootstrap, E: *p* < 0.001, bootstrap).

**Fig. 3:**
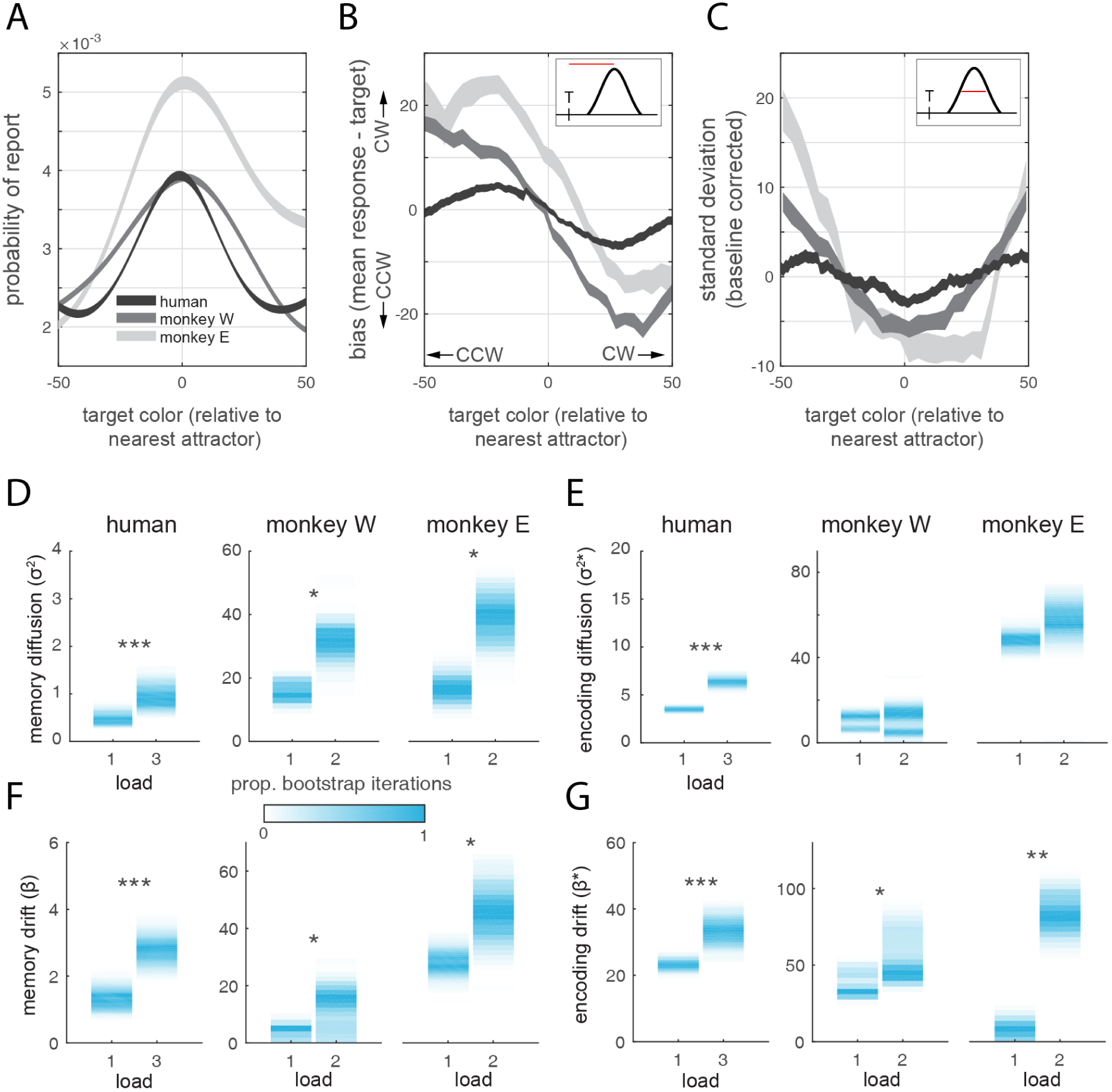
Attractor dynamics explain the frequency, bias, and precision of memory reports, and adapt with load. (A) Probability of report +/- SEM for colors near attractors identified by the dynamical model. (B) Bias +/- SEM around attractors. Inset: bias is calculated as the angular distance between the target and mean report. Positive values indicate clockwise (CW) drift; negative values indicate counter-clockwise (CCW) drift. (C) Standard deviation of the response distribution (inset) +/- SEM around attractors. (D-G) Maximum likelihood parameter fits for the drift and diffusion parameters for human and monkey subjects. Color intensity reflects normalized proportion of bootstrap iterations.

Second, attractors explain bias in working memory reports. Drifting towards an attractor will induce a systematic bias in the reported memory. Indeed, memories counter-clockwise to an attractor location drifted clockwise (and vice-versa, Fig. 3b). The slope of the bias around attractors was significantly negative for both humans (mean slope −0.32 less than zero, *t*(89) = – 7.80, *p* < 0.001, t-test) and monkeys (W: −0.77, *p* < 0.001, bootstrap; E: −0.78, *p* < 0.001, bootstrap).

Model-free analyses showed similar effects. The peaks in the response histogram can be taken as model-free estimates of attractor locations. Aligning the bias around peaks in the response histogram showed a similar pattern (Fig. S5) with a significantly negative slope (H: mean slope −0.42 less than zero, *t*(89) = –7.69, *p* < 0.001, t-test; W: −0.72, *p* < 0.001, bootstrap; E: −0.87, *p* < 0.001, bootstrap). In addition, there was a negative correlation between the slope of the bias and frequency of report (Fig. S6, H: *r*(359) = –0.57, *p* < 0.001; W: *r*(63) = –0.46, *p* < 0.001; E: *r*(63) = –0.18, *p* = 0.155). Note that these results also exclude other possible explanations for a non-uniform response profile, such as subjects guessing with a biased response distribution (Fig. S7).

Third, attractors explain the precision of working memory reports. Memories near an attractor will be more stable: as noise drives the representation away from an attractor, the dynamics will pull it back towards the attractor, resulting in more precise reports with a lower standard deviation (SD, Fig. 2a). Indeed, SD was significantly reduced for colors near attractors (Fig. 3c, H: ∆SD=-2.70, *t*(89) = –4.85, *p* < 0.001; W: −5.46, *p* < 0.001; E: −5.80, *p* < 0.001). Again, model-free analyses supported these conclusions: SD was significantly reduced at the peaks in the response histogram (Fig. S5, H: ∆SD=-3.14, *t*(89) = –2.76*, p* = 0.007; W: −6.14, *p* < 0.001; E: −7.20, *p* < 0.001) and there was a significant correlation between SD and frequency of report (Fig. S6, H: *r*(359) = –0.40, *p* < 0.001; W: *r*(63) = –0.76, *p* < 0.001; E: *r*(63) = –0.86*, p <* 0.001).

These error-correcting properties may be especially critical when memory load is high. High memory load decrease the gain of neural responses (2) which is thought to increase diffusion due to random noise (17). Indeed, the dynamical model showed diffusion during the memory delay increased with memory load (Fig. 3d, 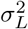, H: *p* < 0.001; W: *p* = 0.021, E: *p* = 0.010) but had a mixed effect during encoding (Fig. 3e, 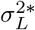, H: *p* < 0.001; W: *p* = 0.459, E: *p* = 0.100). Consistent with the theory that attractor dynamics compensate for diffusion, we saw a commensurate increase in attractor dynamics during the memory delay (Fig. 3f, *β_L_*, H: *p* < 0.001; W: *p* = 0.026, E: *p* = 0.026) and during encoding (Fig. 3g,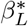, H: *p* = 0.002; W: *p* = 0.024, E: *p* = 0.009). Similar effects were seen in model-free analyses: drift in memory representations, as measured by clustering of memory reports and bias, tended to increase with load (Fig. S1).

While attractors compensate for diffusion, they also induce systematic error into working memory representations. Thus, there is a tradeoff between the finite error caused by drifting toward an attractor and the ever-increasing error associated with diffusion. To test whether attractor dynamics improved overall memory performance, we generated synthetic memory reports from the fitted full dynamical model (‘drift + diffusion’, e.g. Fig. 4a, left) and from a ‘diffusion-only’ model (e.g. Fig. 4a, right, *β* and *β*^*^ = 0). As shown in Fig. 4b, the two models accumulate error at different rates over time. Initially, the mean absolute error is greater in the drift + diffusion model due to memory corruption by drift towards attractor states (*p* < 0.05 for *t* < 9*s*). However, the drift + diffusion model performed significantly better at longer delays (*p* < 0.05 for *t* > 18*s*), with the crossover in performance occurring at *t* = 12*s*, as the attractor dynamics mitigated the effect of diffusive noise over longer periods of time.

**Fig. 4:**
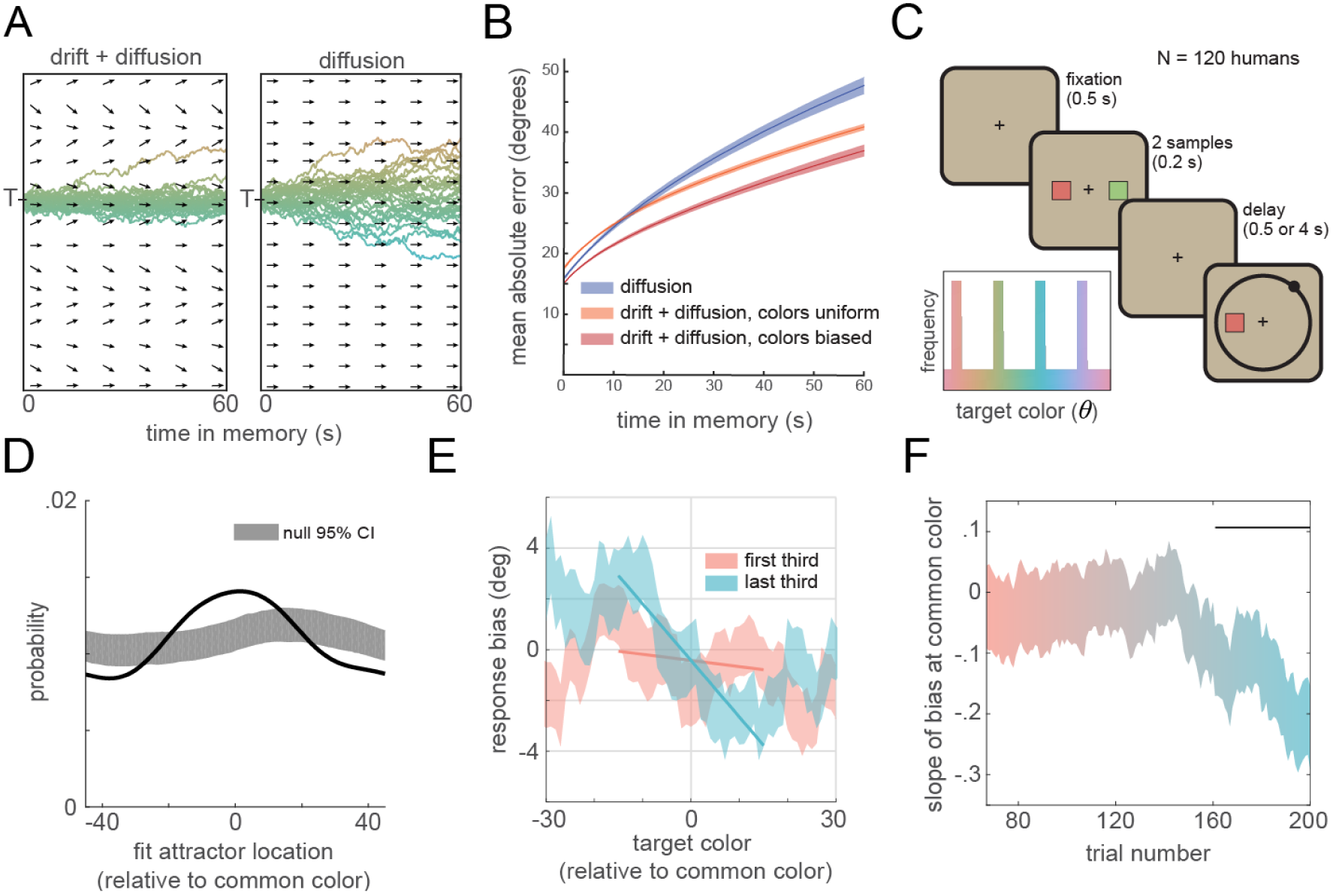
Attractor states reflect the statistics of the environment. (A) Simulated memory trajectories for the full ‘drift + diffusion’ model and the ‘diffusion-only’ model (B) Mean absolute error for the memory of the target color as a function of time for both models. Shaded regions indicate 95% CI. The performance of the diffusion model does not depend on the distribution of target colors. Results are based on the mean parameter fits for the high load condition; similar results were observed for low load. (C) Humans (N=120) performed a color delayed estimation task with two stimuli. Inset: Example color distribution for one subject. Four groups of colors were presented more frequently. Common colors were equally spaced in color space and differed for each subject. (D) Probability of attractor position (estimated from dynamical model fits) relative to common colors (black). Shaded region indicates 95% confidence intervals of the null distribution, based on randomly permuting the location of common colors across subjects. (E) Bias for targets around common colors during the first and last third of trials, with regression lines. (F) Slope of the bias at common colors over trials. Negative values indicate attraction towards common colors. Black line indicates significantly different from zero (*p* < 0.05).

While most laboratory studies (including above) uniformly distribute stimuli across mnemonic space, the distribution of stimuli in the real world is non-uniform (19). When the stimulus distribution is not uniform, the benefit of attractor dynamics on working memory performance is amplified. In particular, if attractors are located at the position of common stimuli, then working memory performance is improved for all *t* (Fig. 4b, red trace). Therefore, in order to minimize working memory errors in a given context, attractor dynamics should change to reflect the statistics of the current environment.

To test whether subjects learned optimal attractor dynamics, a new group of human subjects (N=120) performed a continuous working memory task but with a biased color distribution (Exp. 2, Fig. 4c). During this task, the statistics of the environment were such that half of all stimuli were drawn from one of four common colors (randomly chosen for each subject) while the other half were drawn from a uniform distribution. Both model-free and model-based analyses show participants developed attractors at the common color locations. First, attractors, as identified by fitting the dynamical model, were significantly more likely to appear at the location of common colors than expected by chance (Fig. 4d, *p* < 0.001, randomization test, model fits were limited to trials in which the target color was drawn uniformly). Second, the slope of the response bias around the common colors was significantly less for the last third of trials than the first third (Fig. 4e, *t*(119) = 2.05, *p* = 0.043). This effect grew with experience; the slope around common colors became steeper during the task (Fig. 4f, *r*(133) = –0.747, *p* < 0.001). Finally, subjects were significantly more likely than chance to report common colors, even on the half of trials when the target was drawn from a uniform distribution (Fig. S8, *p* < 0.001, randomization test).

Previous work has emphasized the impact of noise on working memory, arguing this causes a memory to diffuse away from its original state (1, 23, 24). Our results suggest that underlying attractor dynamics have a significant influence on working memory representations. Together, noisy diffusion and drift towards attractors provide a parsimonious explanation of the frequency, bias, and precision of memory reports.

Attractor dynamics are thought to reduce the impact of neural noise in long-term memory (20, 21) and decision making (22, 23) and have been hypothesized to play a similar role in working memory (24–26). Indeed, our results show attractor dynamics within mnemonic space limit the impact of random diffusion, reducing errors in working memory. From an information-theoretic perspective, attractor dynamics effectively compress working memory representations by discretizing the continuous color space. Discretization reduces the information needed to encode the memory, allowing it to be more accurately stored in a noisy system (27, 28). This is consistent with the increase in attractor strength with working memory load; as the information channel becomes crowded with multiple items, compression should increase.

Furthermore, by reflecting the statistics of the world, attractor dynamics act to integrate prior beliefs with noisy stimulus information. In this way attractor dynamics are analogous to Bayesian inference applied over time. At each timestep in memory, drift ‘applies’ the prior to each item in memory (embedded in the attractors), which reflects the posterior of the previous timestep plus random noise. Thus, as time in working memory increases (and stimulus information diffuses), memory representations drift towards prior expectations. This may be a general mechanism for compensating for noise in the brain.

## Materials and Methods

### 1.1 Participants

Thirty-three human subjects participated in Experiment 1 at Princeton University. Seventy-three additional subjects participated in an online version of Experiment 1 via Amazon Mechanical Turk (https://www.mturk.com). One-hundred fifty-five subjects participated in Experiment 2 via Amazon Mechanical Turk. We screened subjects for a minimum of engagement in the task by estimating their probability of random guessing in the task using 3-component mixture model (7). Subjects with an estimated guess rate greater than 20% across all trials were excluded from further analysis, yielding thirty laboratory subjects and sixty online subjects for Experiment 1 and one-hundred twenty online subjects for Experiment 2. This threshold of 20% was set independently based on analysis of a separate pilot cohort of online subjects (N = 57). Subjects recruited online via Mechanical Turk have previously been used to study working memory and have performance comparable to lab subjects (29, 30). We observe similar qualitative behavior between online and lab subjects (Fig. S9) and report their behavior together in the main text. All subjects attested that they had normal or corrected-to-normal vision. We confirmed that subjects had normal color vision using the Ishihara Color Blindness Test. Subjects provided informed consent in accordance with the Princeton University Institutional Review Board.

Two adult male rhesus macaques (8.9 and 12.1 kg) performed the Experiment 1 in accordance with the policies and procedures of the Princeton University Institutional Animal Care and Use Committee.

### 1.2 Experiments

#### 1.2.1 Experiment 1 - humans

For the laboratory version of Experiment 1 we presented stimuli on a CRT monitor positioned at a viewing distance of 60 cm. We calibrated the monitor using an X-Rite i1Display Pro colorimeter to ensure accurate color rendering. During the experiment, participants were asked to remember the color and spatial location of either 1 or 3 square “sample” stimuli. The color of each sample was drawn from 360 evenly spaced points along an isoluminant circle in CIE L*a*b* color space. This circle was centered at (L = 60, a = 22, b = 14) and the radius was 52 units. Colors were drawn pseudorandomly, with the caveat that colors presented on the same trial had to be at least 22° apart in color space. The samples measured 2° of visual angle (DVA) on each side. Each sample could appear at one of eight possible spatial locations. All possible locations had an eccentricity of 4.5 DVA and were positioned at equally spaced angles relative to central fixation (0, 45, 90, 135, and 180° clockwise and counterclockwise relative to the vertical meridian). The dimensions of the stimuli for the online experiment were defined by pixels rather than degrees of of visual angle. The samples had an edge length of 30 pixels and were presented at an eccentricity of 170 pixels.

Participants initiated each trial by clicking the mouse and by fixating a cross at the center of the screen (Fig. 1a). After 500 ms of fixation, one or three samples (the “load”) appeared on the screen. The samples were displayed for 200 ms and then were removed from the screen. Participants then experienced a memory delay of 1 second or 7 seconds, after which a response screen appeared. The response screen consisted of the outline of a square at one of the previous sample locations (the “probe sample”) and a response interface consisting of a circle on a ring. Participants used the mouse to drag the circle around the ring, which changed the color of the probe sample. The angular position of the circle on the ring corresponded to a particular angle in color space. The mapping between circle position and color space was randomly rotated on each trial to minimize contributions from spatial memory. We instructed participants to adjust the color of the probe sample to match the color of the sample that had previously appeared at that location as closely as possible. We told participants that accuracy was more important than speed but that they should respond within a few seconds. There was no time limit on the response. All human participants completed 200 trials.

We monitored the eye position of the lab participants using an Eyelink 1000 Plus eyetracking system (SR Research). Participants had to maintain their gaze within a 2° circle around the central cross during initial fixation and sample presentation, or else the trial was aborted and excluded from analysis.

#### 1.2.2 Experiment 1 - monkeys

We presented stimuli on a Dell U2413 LCD monitor optimized for color rendering. The monitor was positioned at a viewing distance of 58 cm. We calibrated the monitor using an X-Rite i1Display Pro colorimeter to ensure accurate color rendering. Sample colors were drawn from 64 evenly spaced points along an isoluminant circle in CIE L*a*b* color space. This circle was centered at (L = 60, a = 6, b = 14) and the radius was 57 units. Slightly different color wheels were used for the humans and the monkeys to accommodate the gamut of the different monitors used in each experiment. Nevertheless, colors corresponding to the same angle in each color wheel are extremely similar in appearance. The edges of the samples measured 2° of visual angle. Each sample could appear at one of two possible spatial locations: at 5 DVA eccentricity from fixation and 45° clockwise and counterclockwise from the horizontal meridian.

We adapted Experiment 1 so that it could be performed by non-human primates. The animals initiated each trial by fixating a cross at the center of the screen. After 500 ms of fixation, one or two samples appeared on the screen. The samples were displayed for 500 ms, followed by a memory delay of 500 ms or 1500 ms. Next, a symbolic cue was presented at fixation for 300 ms. This cue indicated which sample (top or bottom) the animal should report in order to get juice reward. The response screen consisted of a ring 2° thick with an outer radius of 5°. The animals made their response by breaking fixation and saccading to the section of the color wheel corresponding to their report. This ring was randomly rotated on each trial to prevent motor planning or spatial encoding of memories. The animals received a graded juice reward that depended on the accuracy of their response. The number of drops of juice awarded for a response was determined according a circular normal (von mises) distribution centered at 0° error with a standard deviation of 22°. This distribution was scaled to have a peak amplitude of 12, and non-integer values were rounded up. When response error was greater than 60°, no juice was awarded and the animal experienced a short time-out of 1 to 2 seconds. Responses had to be made within 8 seconds; in practice, this restriction was unnecessary as response times were on the order of 200-300 ms. We analyzed all completed trials (trials on which the animal successfully maintained fixation and saccaded to the color wheel, regardless accuracy). Monkey W completed 15,787 trials over 26 sessions and Monkey E completed 16,601 trials over 17 sessions.

We monitored the eye position of the animals using an Eyelink 1000 Plus eyetracking system (SR Research). The animals had to maintain their gaze within a 2° circle around the central cross during the entire trial until the response, or else the trial was aborted and the animal received a brief timeout. Trials during which the animal broke fixation were excluded from analysis.

#### 1.2.3 Experiment 2

The stimuli and procedures for Experiment 2 (Fig. 4c) were similar to those for the online version of Experiment 1. We shortened the memory delays to 500 ms and 4000 ms to reduce the length of the experiment. Participants saw 2 samples on each trial. Critically, the color of the samples was no longer always drawn uniformly from the circular color space. Rather, for each sample, there was a 50% chance that the color of that sample would be drawn from a biased distribution (Fig. 4c). This biased distribution consisted of four equally-spaced clusters of common colors. Each cluster was 20° in width. Each participant was exposed to a unique set of common colors as the cluster means were shifted by a single random phase for each subject.

### 1.3 Analysis

#### 1.3.1 Effects of load and time on mean error

We analyzed mean absolute error for human subjects using a 2×2 repeated measures ANOVA with factors load, delay time, and their interaction. We analyzed each monkey’s data by fitting the equivalent regression model to their mean error in each condition. We obtained bootstrapped confidence intervals for each regression coefficient by re-sampling trials with replacement from each monkey’s dataset and refitting the regression model on each iteration (1000 iterations). We also used this method to analyze the effect of load and time on clustering and mean bias (**Fig. ED1**).

#### 1.3.2 Clustering Metric

We observed that the distribution of reported hues *θ̂* are clustered relative to the distribution of target hues Θ. To quantify this phenomenon, we developed a simple clustering metric. This metric relies on the fact that entropy is maximized for uniform probability distributions. In contrast, probability distributions with prominent peaks will have lower entropy. Because the target hues are drawn from a circular uniform distribution, the entropy of the targets *H*(Θ) will be relatively high. If that subject’s responses are clustered, their entropy *H*(*θ̂*) will be relatively low. Taking the difference of these two values yields a clustering metric *C*. Negative values of *C* suggest greater clustering:

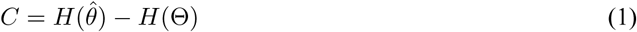

where:

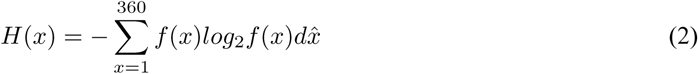

We estimated the pdf of the responses *f* (*θ̂*) and the targets *f* (Θ) using kernal density estimation (Matlab CircStat toolbox, kernal width = 10°). Our goal was to quantify the clustering of reports for items in memory. Random guesses (6, 7) confound this analysis by contributing a uniform component to the response distribution that varies systematically as a function of load and time. To address this, we estimated the proportion of responses due to guessing using mixture models (6, 7) and removed a uniform component from the response distribution *f* (*θ̂*) and the target distribution *f* (*θ*) equal in area to the guess rate and then renormalized each.

#### 1.3.3 Bias and standard deviation of memory reports

To dissociate systematic and unsystematic sources of error in memory, we calculated the bias and standard deviation of memory reports across color space (20° bins). Bias refers to the distance between the the target color and the mean reported color. We calculated the slope of bias around negative-slope zero-crossings in each subject’s fit drift function (Experiment 1), around significant peaks in each subject’s response histograms (Experiment 1), and around commonly presented presented colors (Experiment 2) by fitting a line to the bias +/- 15° around the point of interest. For monkey subjects, we boostrapped confidence intervals for slope and standard deviation by resampling trials with replacement. To compute the bias and SD for the non-uniform guessing strategy (**Fig. ED7**, dashed lines), we performed 1000 iterations of a randomization test where memory reports were shuffled with respect to the target colors and report the mean bias and SD for each target color across iterations.

To identify significant peaks in subjects’ response histograms (Experiment 1), we first estimated the PDF of subjects’ responses using kernal density estimation. We identified possible peaks as samples larger than their two neighboring samples and recorded their amplitude. We then repeated this analysis on the distribution of targets, resampling with replacement to create a null distribution of peak amplitudes. Peaks in the original response distribution with an amplitude greater than the 95th percentile relative to the null were deemed significant. We identified negative-slope zero-crossings in the fit drift function of each subject by identifying peaks in the numerical integral of the drift function. Peaks with a prominence in the 20th percentile or lower across subjects were excluded from analysis.

#### 1.3.4 Dynamic Model

We developed a quantitative model to describe how items in memory change over time. We assume that two distinct influences may make memory dynamic. First, systematic biases may cause memories to “drift” towards stable attractor states over time. Second, memories may be perturbed by unsystematic random noise. We model memory using a stochastic ordinary differential equation that captures both of these influences:

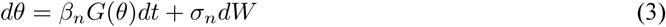

This equation describes the time evolution of a color memory *θ* (a circular variable corresponding to an angle in our circular color space) under the influence of some deterministic dynamics defined by *G* (the “drift”) as well as an additive white noise process *W* with variance *σ*^2^. *β_n_* sets the gain of the drift. Thus, *β_n_G*(*θ*)*dt* describes influence of drift and *σ_n_dW* the influence of random noise on memory. To test the hypothesis that memory load influences these dynamics we fit a separate *β* and *σ* for each load *n*.

Based on the clustering we observe in the data, it seems likely that *G*(*θ*) is a nonlinear function. We needed a relatively parsimonious way of describing *G*(*θ*) that still gave us enough flexibility to describe this nonlinearity. So, for each subject, we defined *G*(*θ*) using a basis set consisting of twelve first derivatives of the von mises distribution separated by 1 standard deviation on the interval (0, 2*π*):

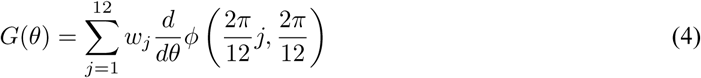

where *ϕ* is a von mises distribution parameterized by a mean and standard deviation. We then divided *G*(*θ*) by its maximum absolute value. This normalization procedure aids the interpretation of *β*: it is the maximum instantaneous drift rate. We decided to use 12 basis functions in a principled fashion using AIC.

To fit the model described in equation 3 to subject data, we needed to describe the time evolution of *θ* probabilistically. So, we rewrote equation 3 as a Fokker-Planck equation, a partial differential equation that tracks probability density function of *θ* over time:

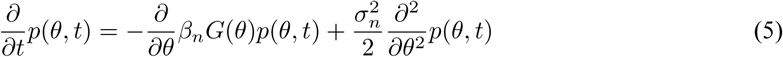

In order to track probability mass, we discretized our 1-dimensional state space (the value of *θ*) into 100 evenly spaced bins from 1° to 360°. Once discretized, the change in *p*(*θ*, t) over a given timestep *dt* can be described by a Markov transition matrix *M_n_*:

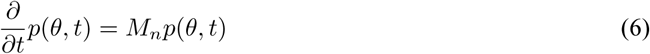

This discretized approximation can be solved analytically in time, yielding:

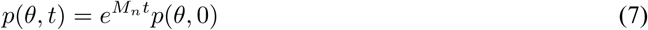

where *p*(*θ*, 0) is the initial state of memory after encoding.

We wanted to dissociate load-driven changes in the dynamics of memory and encoding. To capture differences in encoding, we allowed the state of a memory at the start of the delay, *p*(*θ*, 0), to vary as a function of load. To simulate the encoding process, we first initialized a narrow probability density *P*_0_(Θ) that reflects the color of the target stimulus. *P*_0_ is a von mises distribution with mean equal to the target color Θ and a standard deviation of 0.1 radians:

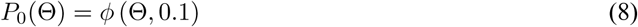

We then allowed *P*_0_ to propogate for a 1 second “encoding period” according to the following differential equation:

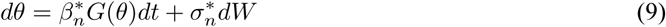
where 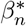 and 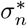 interact to set the bias and variance of the encoded memory. Therefore, *p*(*θ*, 0) is calculated as:

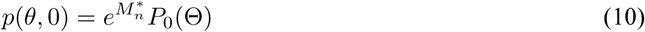

and the final probability distribution describing the memory of the target hue after a memory delay of t seconds on a trial with load *n* is:

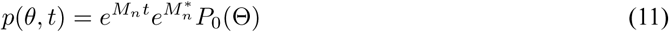

Equation 11 describes the probability distribution for the memory of the target color Θ at time *t*. However, our goal is to predict the subject’s *report* on a particular trial, *p*(*θ̂*, *t*), which does not just depend on the color of the target (6, 7). On some trials, subjects may erroneously report their memory of one of the non-target samples, 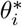. To account for these “swap errors”, we also allow the memory of any distractor colors 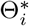 to evolve according to equation 11. On other trials, subjects may experience complete failures of memory, resulting in random guessing. To account for these additional influences, we estimated each subject’s probability of committing swap errors and guessing, and, for each trial, computed a mixture of the target memory distribution, the non-target memory distributions, and a uniform component:

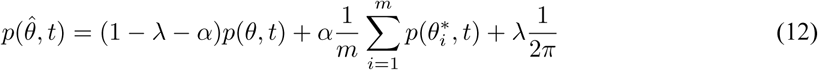

where *m* is the number of non-target colors (0 or 2 for humans, 0 or 1 for monkeys). *α* and *λ* represent the probability of swap errors and guesses, respectively. They are linear functions of *t* parameterized by a slope *a* and intercept *b*. We estimated a unique lambda function for each load. We found the maximum likelihood estimate (joint likelihood across trials) of the free parameters (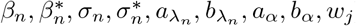) for each subject using gradient descent in Matlab. To obtained boostrapped distributions of the parameter distributions for human subjects, we repeatedly resampled the parameters fit to each subject with replacement and took the mean of these values. To obtain bootstrapped distributions for monkey subjects, we repeatedly resampled each monkey’s pool of trials with replacement and repeated the fitting process.

Model fits indicated that random guessing increased with time for human subjects (Fig. S10), consistent with previous literature (3-5). Guessing decreased with delay, however, for the two monkeys. We wanted to ensure that tradeoffs between guessing and other parameters, such as the rate of diffusion, were not driving the effects of increased drift and diffusion with load. So, we fit different versions of the model in which we systematically simplified our parameterization of guess rate. Across the two monkeys, model comparison using AIC and BIC indicated that the full model was the best fit to the data. Regardless, for all models, drift and diffusion increased with load, indicating that this is a stable feature (Section).

#### 1.3.5 Simulated performance of the drift + diffusion and diffusion models

We wanted to identify if attractor dynamics might be normative and enhance the fidelity of memory. To do this, we computed the expected mean error for the memory of a target color as a function of delay time for the full dynamic model with attractor dynamics (“drift + diffusion”) and a model without attractor dynamics (“diffusion”). The drift and diffusion parameters of the drift + diffusion model were set to the mean fit parameters for the human subjects in Experiment 1. The parameters of the diffusion model were identical except that *β_n_* and 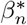 were set to zero. To create a representative drift function, we fit our basis set to the numerical derivative of the PDF of the response distribution for human subjects (normalized to have a maximum absolute value of one), which yields attractors at locations in color space where they are most frequently observed (i.e., at commonly reported colors). To create a biased target distribution, we integrated the drift function, substracted the minimum of the result to avoid negative values, and normalized the result to have an area of one. Finally, we computed the mean expected error for each model and timepoint. Confidence intervals were obtained by repeating this procedure 1000 times, resampling subjects with replacement.

## Acknowledgements

We thank A. Piet for helpful discussions, B. Morea and H. Weinberg-Wolf for assistance with NHPs, and F. Bouchacourt, A. Libby, and P. Kollias for comments. This work was supported by NIMH R56MH115042 and ONR N000141410681 to TJB, an NDSEG fellowship to MFP, and McKnight Foundation, Simons Collaboration on the Global Brain (SCGB AWD1004351) and the NSF CAREER Award (IIS-1150186) to JWP.

## Author contributions

MFP and TJB conceived of the experiments; MFP, BD, JWP, and TJB designed the dynamical model; MFP and BD implemented the model; MFP collected and analyzed the data; MFP and TJB wrote the original draft; MFP, BD, JWP, and TJB discussed the results and prepared the final draft.

## Author information

Authors declare no competing interests. Correspondence and requests for materials should be addressed to TJB.

## Data availability

All data that support the findings of this study are available from the corresponding author upon request.

---

**Figure S1:**
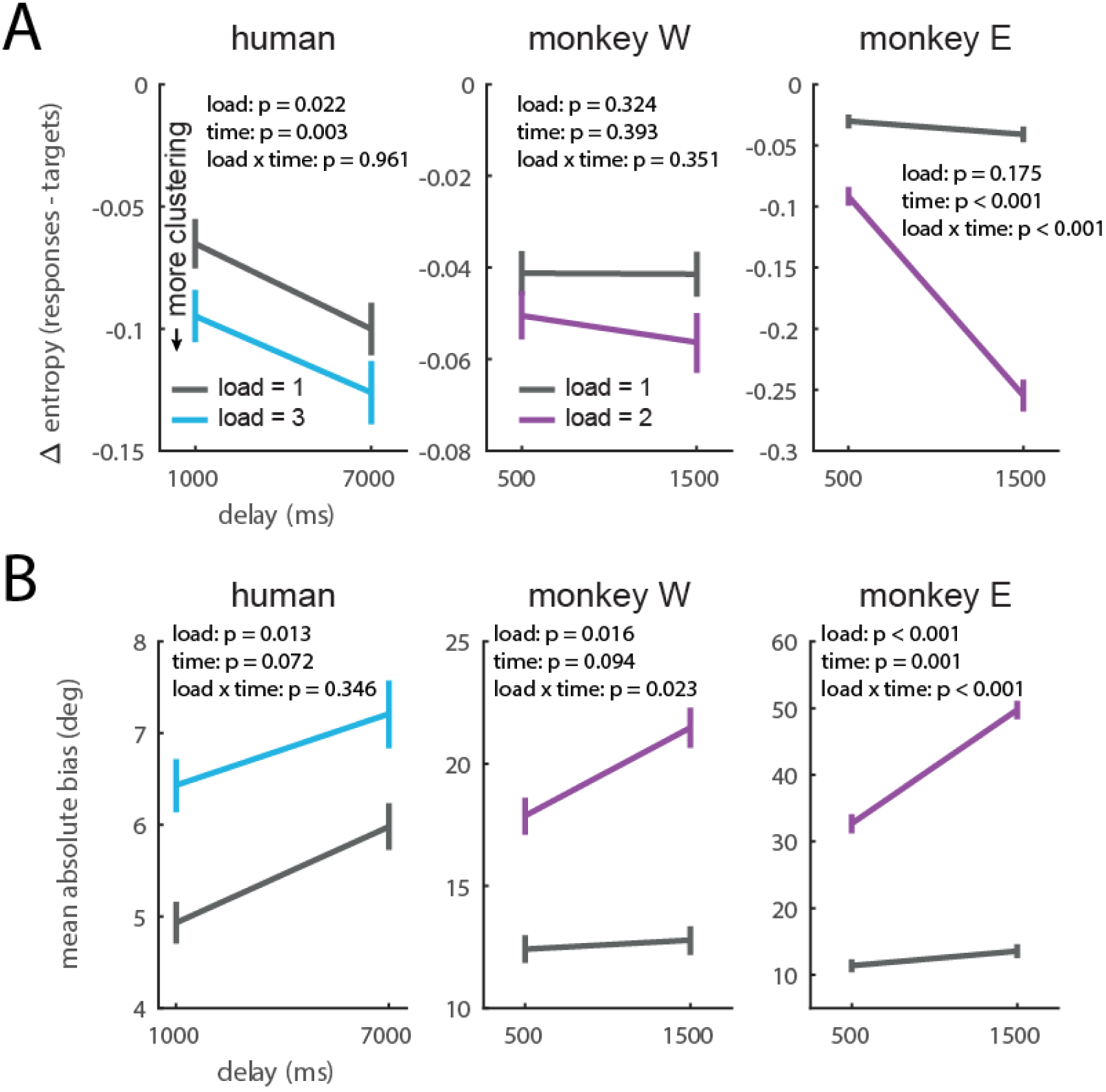
(A) Difference in entropy between the response distribution and target distribution for humans and monkeys as a function of load and delay. More negative values indicate more clustered memory reports. (B) Mean absolute bias (averaged across all target colors) for humans and monkeys as a function of load and delay.

**Figure S2:**
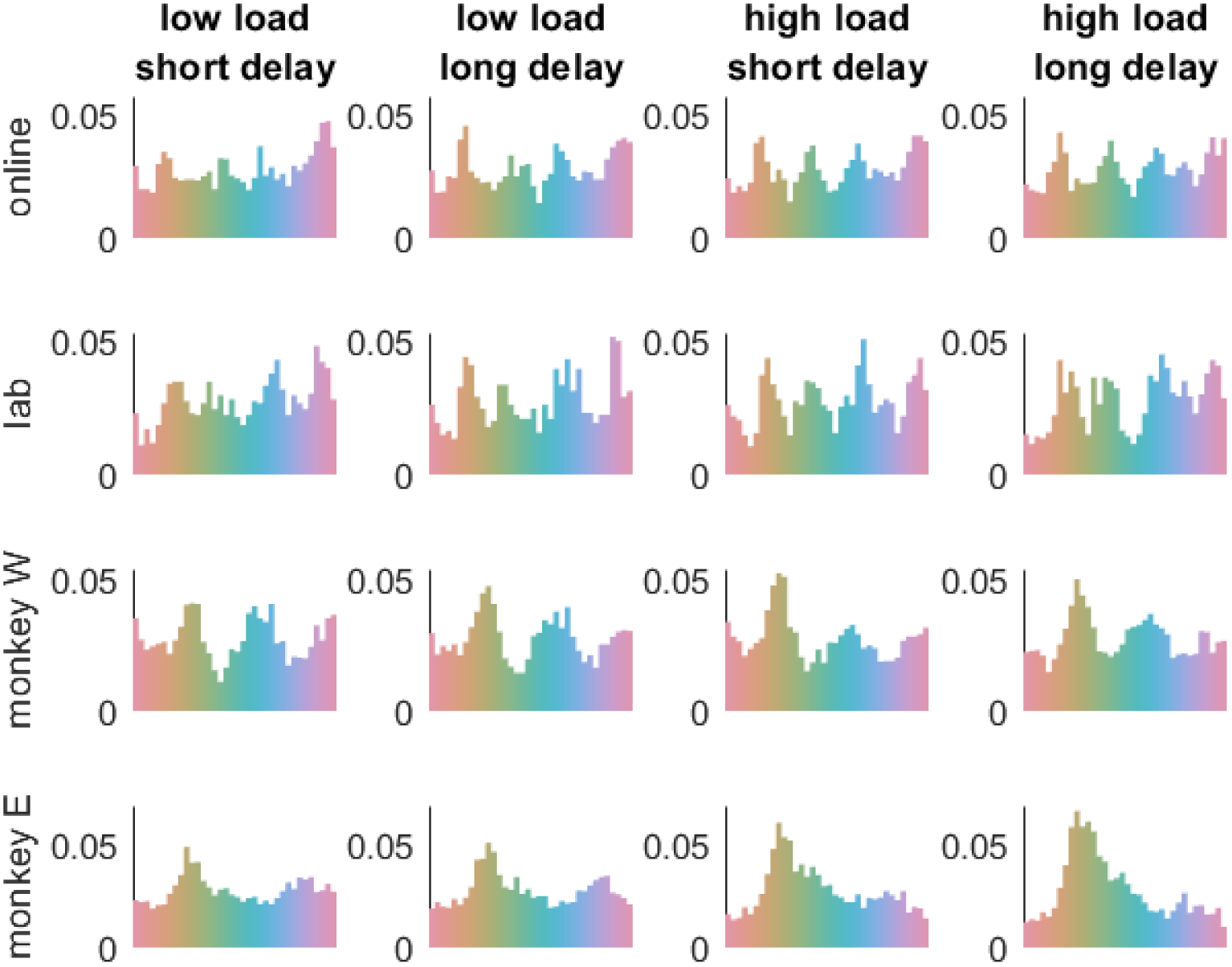
Response histograms for humans and monkeys by condition.

**Figure S3:**
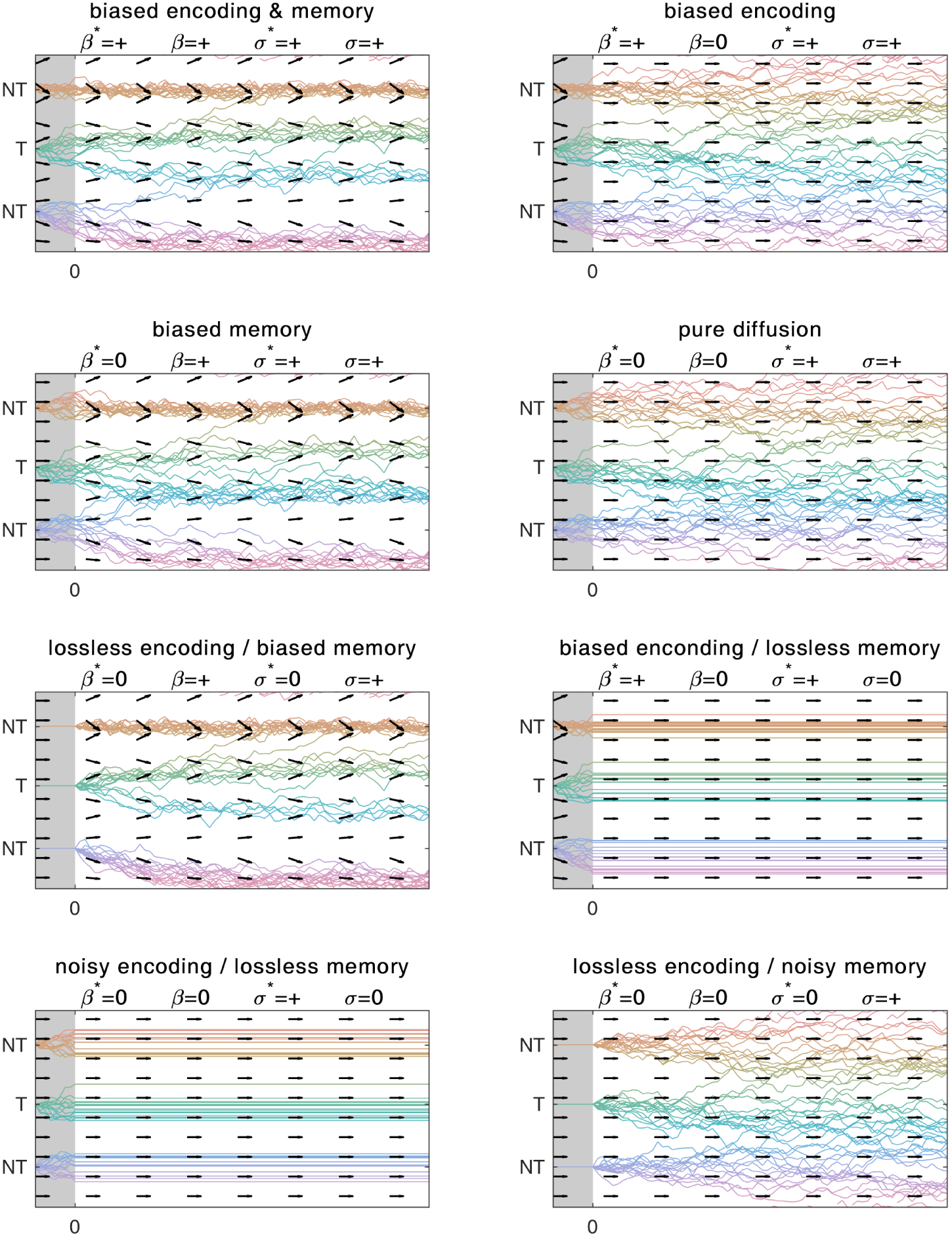
The model can display attractor dynamics or pure diffusion during encoding and memory. Different model behaviors are presented categorically but behaviors along a continuum between these states are of course possible. Further dimensionality is added by the fact that drift and diffusion parameters can increase, decrease, or remain unchanged with increasing load.

**Figure S4:**
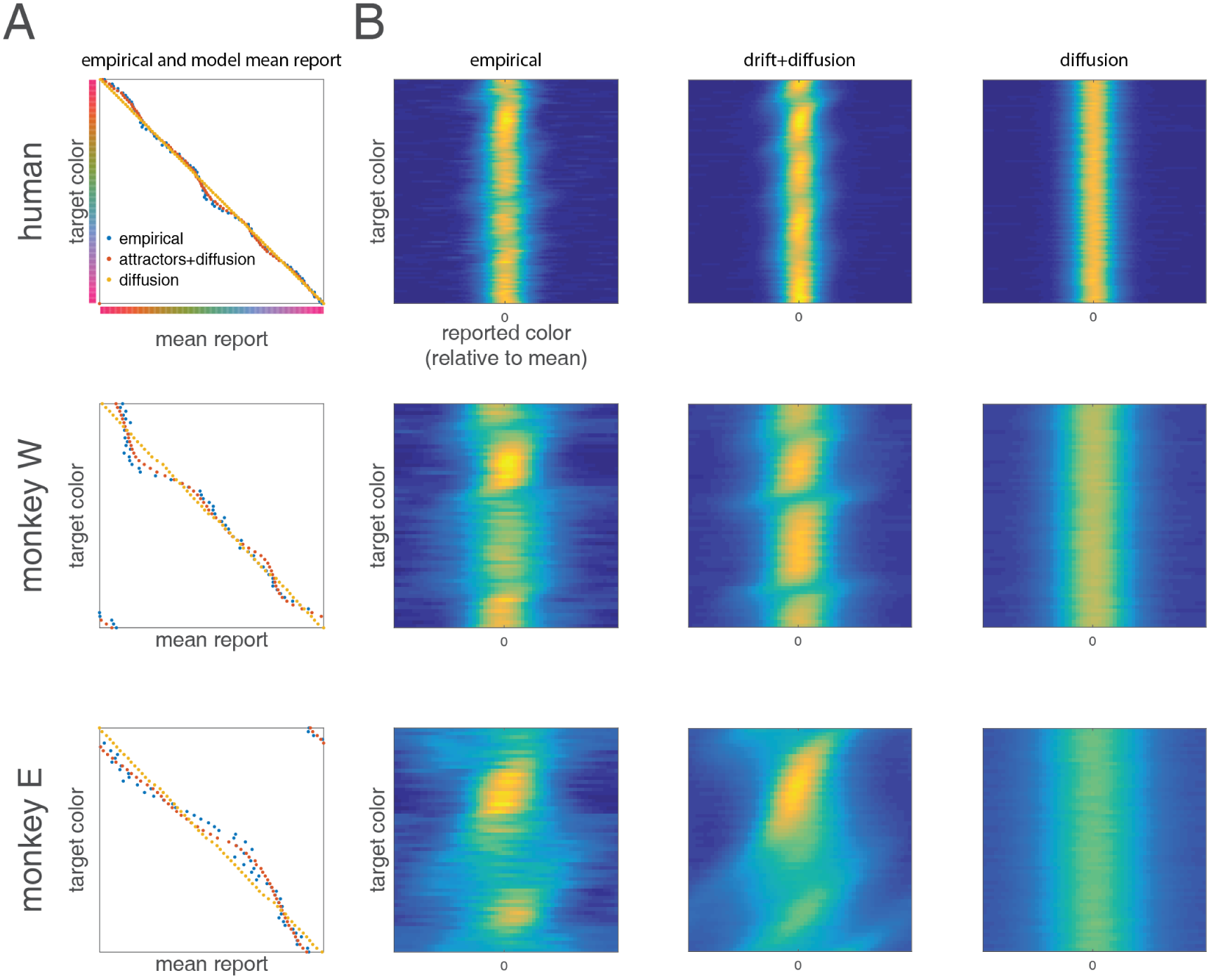
The full model (drift+diffusion) captures more of the variance in responses (Fig. 2) due to its ability to capture both (A) variance in bias across color space and (B) variance in precision across color space.

**Figure S5:**
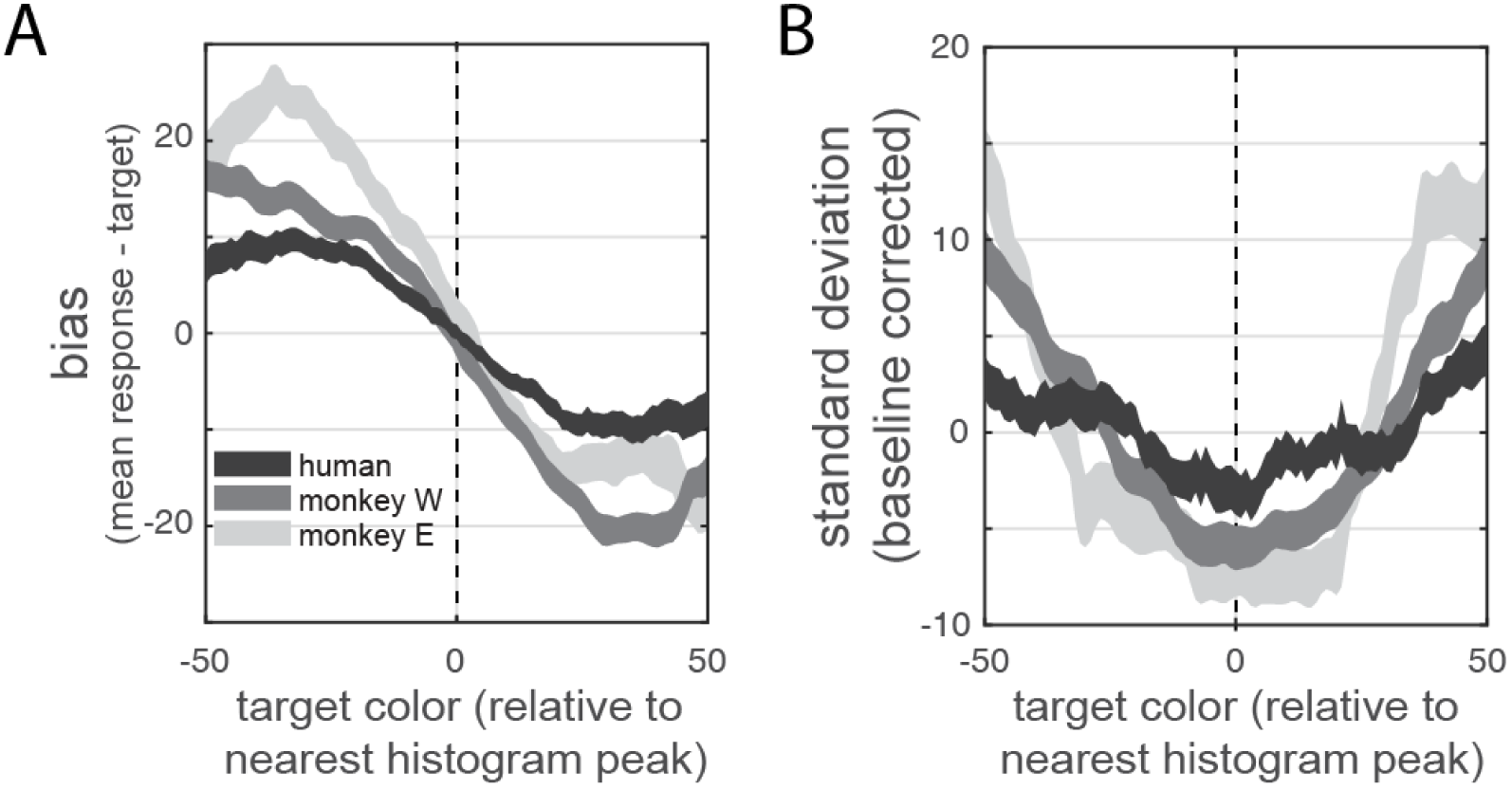
(A) bias and (B) standard deviation of memory reports around putative attractors, identified as significant peaks in subjects’ distribution of reported colors.

**Figure S6:**
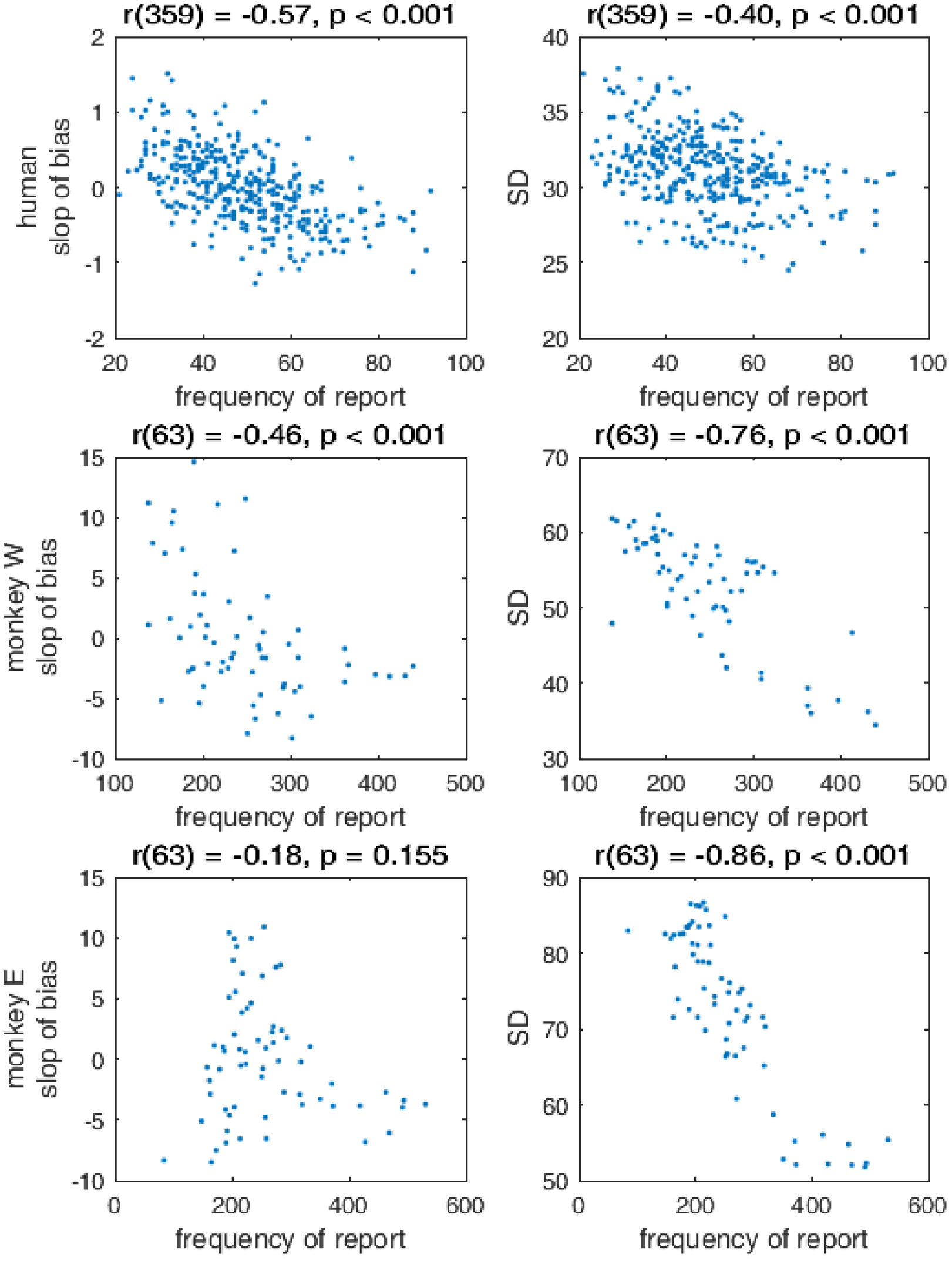
Left: correlation between slope of bias and frequency of report across color space. Right: correlation between standard deviation and frequency of report across color space. Note that the degrees of freedom differ because humans were presented with 360 unique target colors and monkeys were presented with 64.

**Figure S7:**
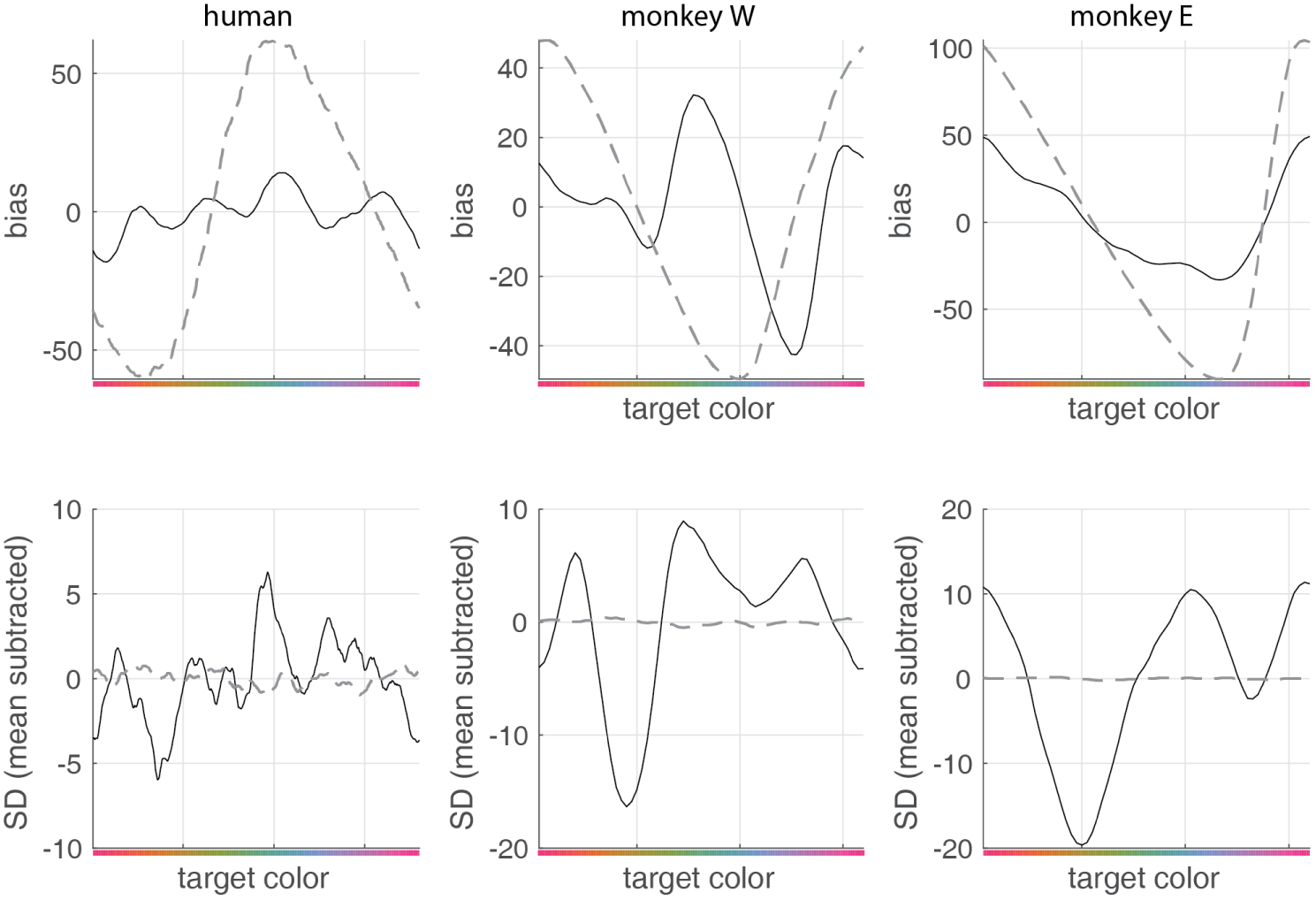
Bias and SD as a function of target color for humans and both monkeys (black lines). Gray dashed lines indicate the predicted bias and SD for a non-uniform guessing strategy, in which the probability of a subject reporting one of the frequently-reported colors does not depend on the identity of the target (see Methods).Non-uniform guessing provides a poor account of the pattern of bias and guessing across target colors. Under non-uniform guessing, bias depends only on the distance between the taget color and the mean reported color across all trials (top row). Additionally, SD does not vary significantly as a function of target color (bottom row).

**Figure S8:**
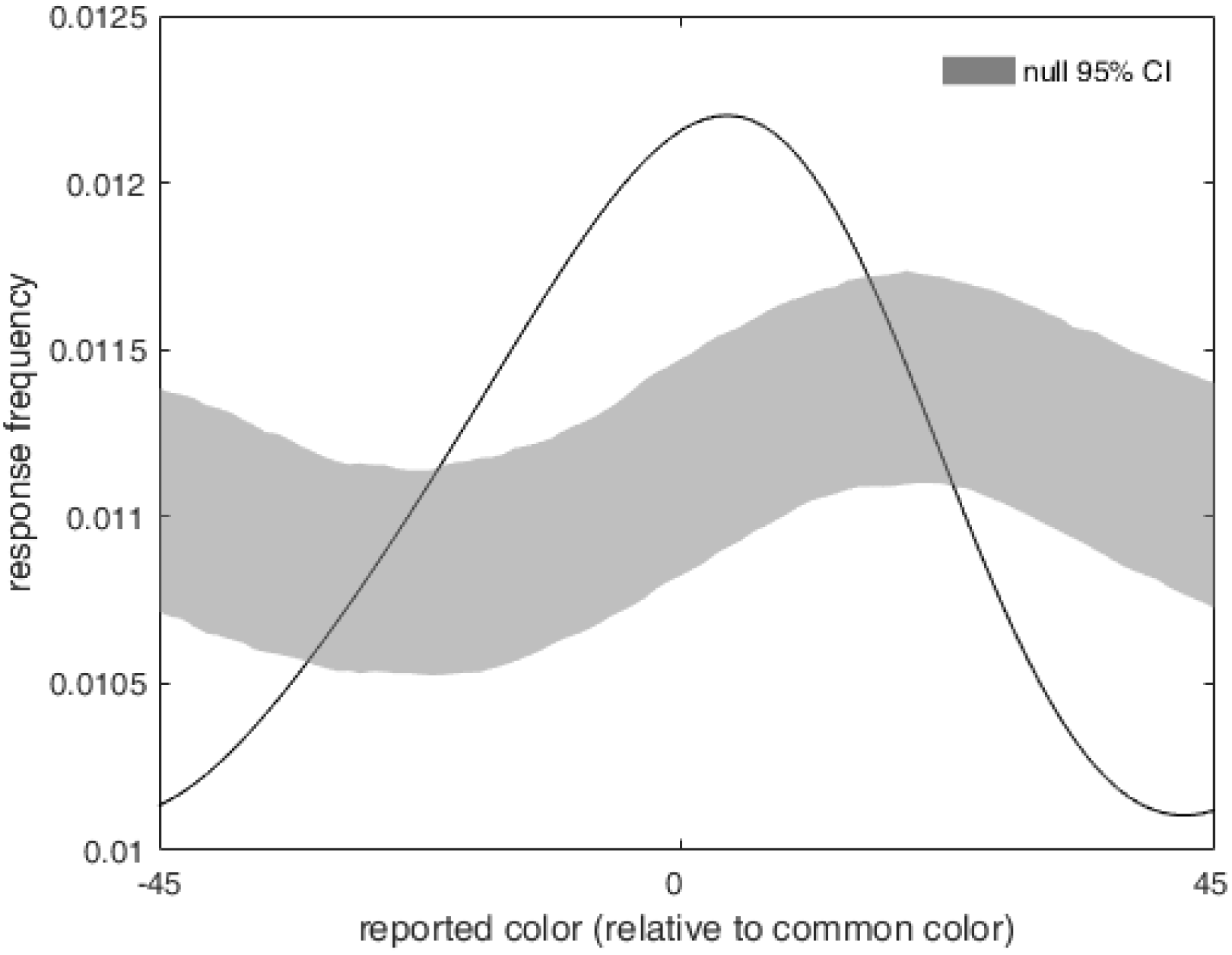
Probability of report as a function of distance from common colors on trials with targets drawn from a uniform distribution.

**Figure S9:**
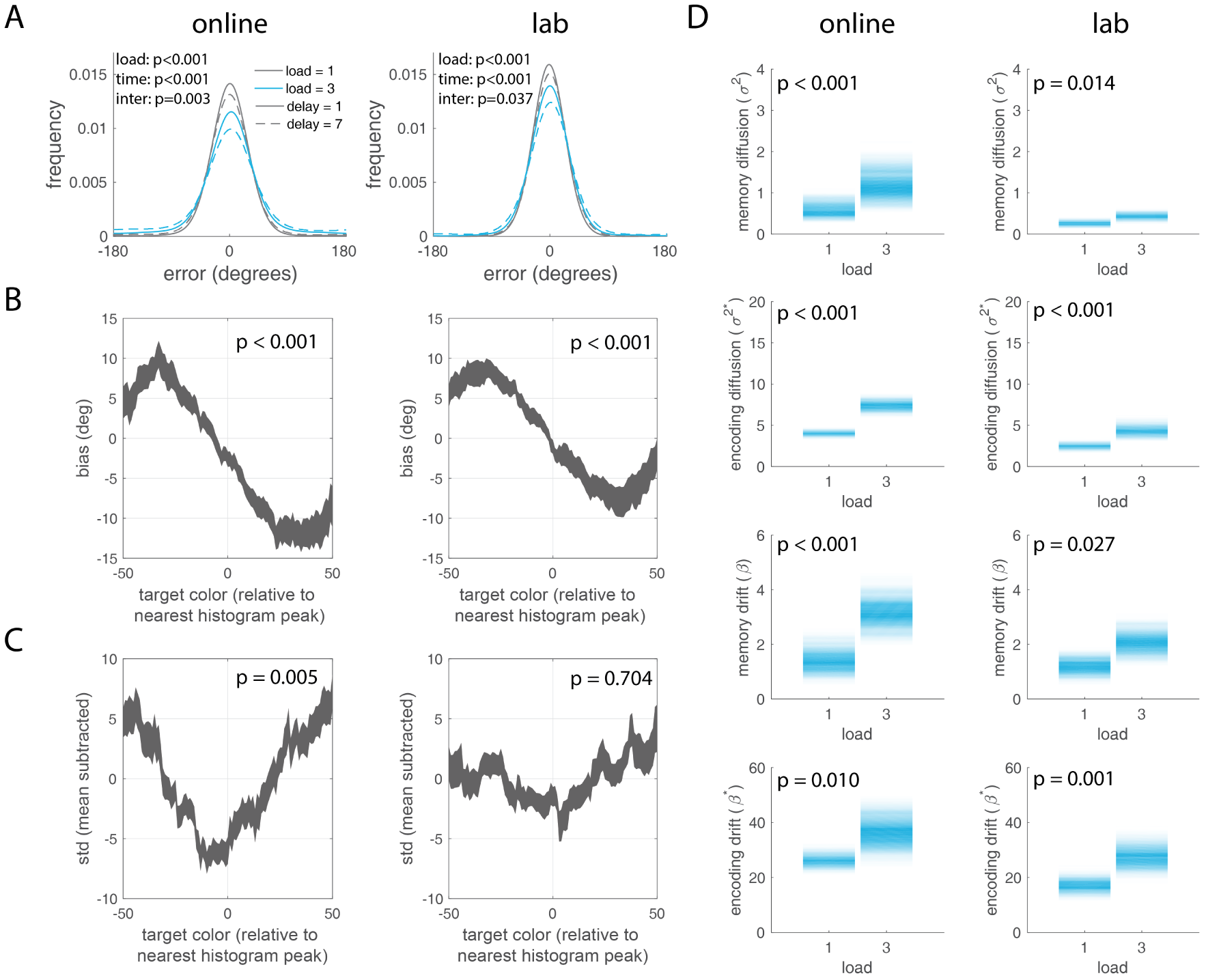
Online and lab subjects show qualitatively similar behavior. P-values reflect (A) the results of a repeated-measures ANOVA predicting mean error as a function of load and time, as in text describing Fig. 1b; (B) a t-test of the slope of bias at histogram peaks vs zero, as in text describing Fig. ED5; (C) a t-test of the relative standard deviation of memory reports at histogram peaks vs zero, as in text describing Fig. ED5; (D) The effect of set size on the diffusion and drift parameter fits from the dynamic model, as in the text describing Fig. 3d-g.

**Figure S10:**
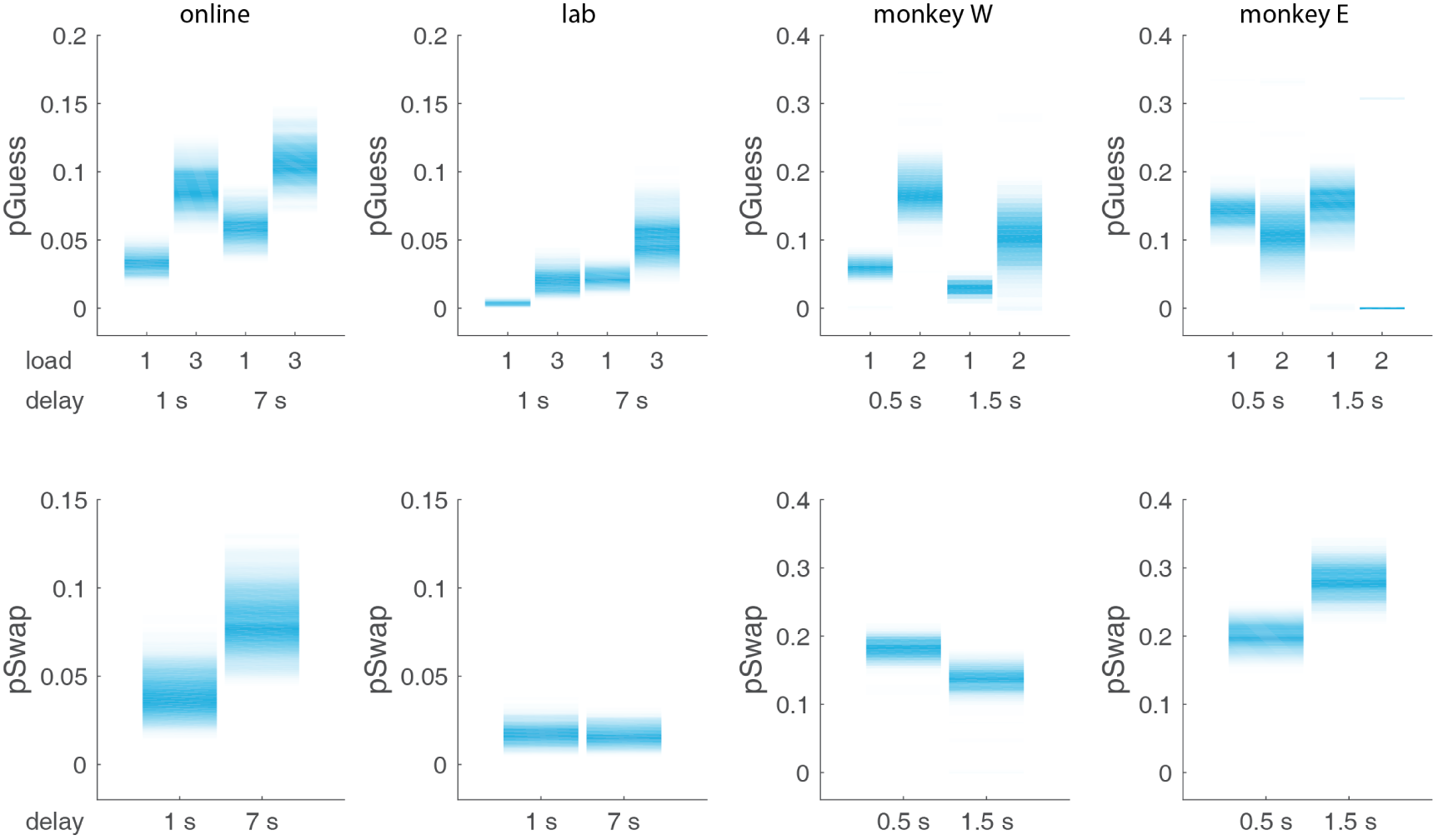
Estimated guess and swap probabilities from dynamic model fits for each load and delay. Color intensity reflects normalized proportion of bootstrap iterations.

**Table S1:**
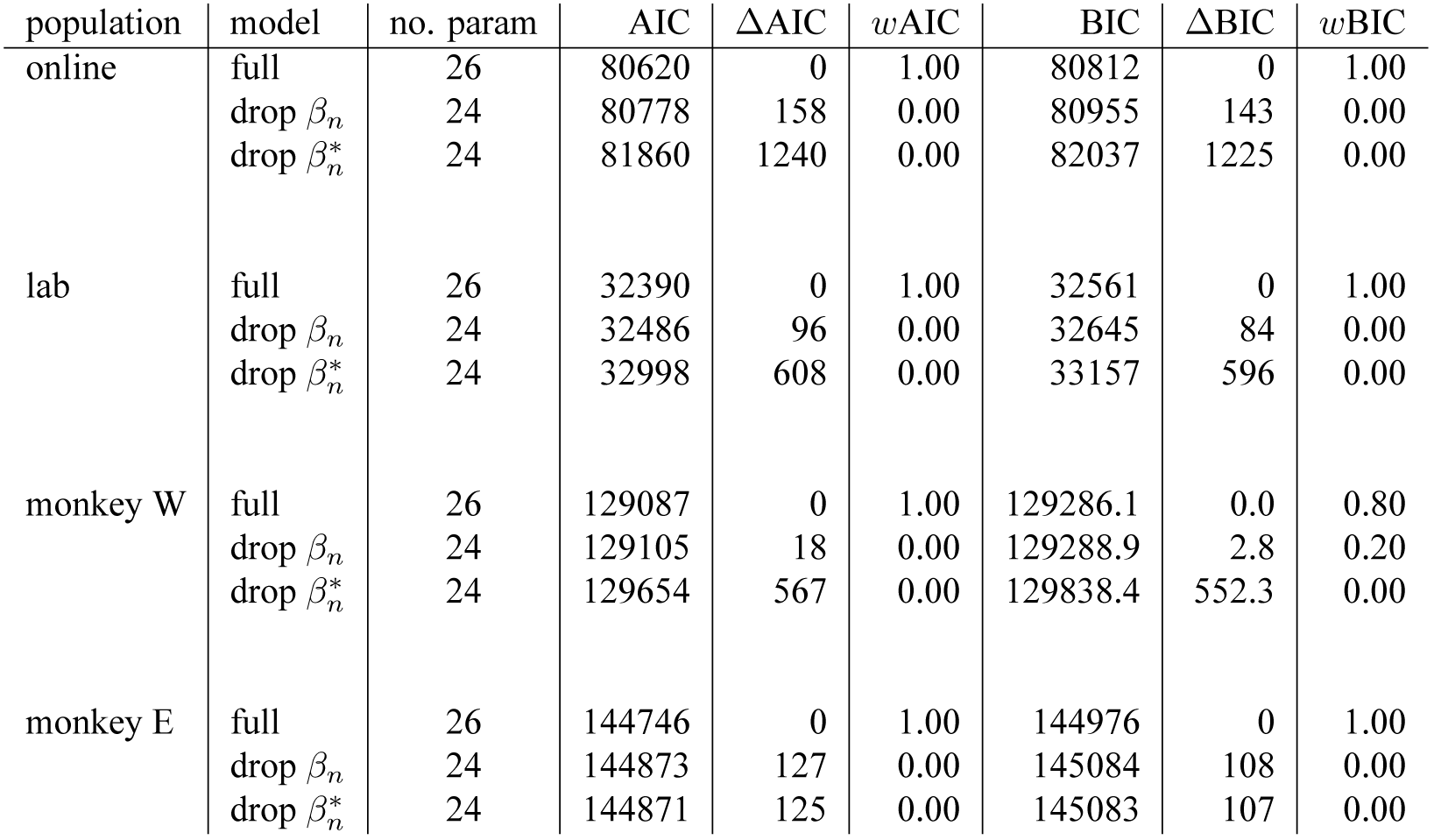
AIC and BIC values for full model and competing models without attractor dynamics during encoding or maintenance. *w*AIC and *w*BIC values indicate the probability that the given model is the best model in the set.

| population | model | no. param | AIC | ΔAIC | $w$AIC | BIC | ΔBIC | $w$BIC |
| --- | --- | --- | --- | --- | --- | --- | --- | --- |
| online | full | 26 | 80620 | 0 | 1.00 | 80812 | 0 | 1.00 |
| | drop $\beta_n$ | 24 | 80778 | 158 | 0.00 | 80955 | 143 | 0.00 |
| | drop $\beta_n^*$ | 24 | 81860 | 1240 | 0.00 | 82037 | 1225 | 0.00 |
| lab | full | 26 | 32390 | 0 | 1.00 | 32561 | 0 | 1.00 |
| | drop $\beta_n$ | 24 | 32486 | 96 | 0.00 | 32645 | 84 | 0.00 |
| | drop $\beta_n^*$ | 24 | 32998 | 608 | 0.00 | 33157 | 596 | 0.00 |
| monkey W | full | 26 | 129087 | 0 | 1.00 | 129286.1 | 0.0 | 0.80 |
| | drop $\beta_n$ | 24 | 129105 | 18 | 0.00 | 129288.9 | 2.8 | 0.20 |
| | drop $\beta_n^*$ | 24 | 129654 | 567 | 0.00 | 129838.4 | 552.3 | 0.00 |
| monkey E | full | 26 | 144746 | 0 | 1.00 | 144976 | 0 | 1.00 |
| | drop $\beta_n$ | 24 | 144873 | 127 | 0.00 | 145084 | 108 | 0.00 |
| | drop $\beta_n^*$ | 24 | 144871 | 125 | 0.00 | 145083 | 107 | 0.00 |

**Table S2:** Maximum likelihood estimates for drift and diffusion parameters for models with different parameterizations of guessing probability. An ‘x’ indicates that a parameter is included in a given model. For the most flexible model (model 1, identical to that reported in the main text), guessing is effectively parameterized by a constant term C, a coefficient determining an effect of load on guessing (L), a coefficient determining an effect of memory delay on guessing (D), and an interaction term (I). Successive models drop combinations of these terms, yielding less flexibility in how guessing changes with load and time. For example, for model 5, p(Guess) is constant across load and time. Regardless of the parameterization, however, drift and diffusion consistently increase with load during both encoding and memory.

| subject | model | p(Guess) eq. |  |  |  | parameter MLE |  |  |  |  |  |  |  |
| --- | --- | --- | --- | --- | --- | --- | --- | --- | --- | --- | --- | --- | --- |
| | | C | L | D | I | $\sigma_1$ | $\sigma_2$ | $\sigma_1^*$ | $\sigma_2^*$ | $\beta_1$ | $\beta_2$ | $\beta_1^*$ | $\beta_2^*$ |
| monkey W | 1 | x | x | x | x | 15 | 31 | 12 | 13 | 4 | 15 | 33 | 45 |
|  | 2 | x | x | x |  | 15 | 29 | 12 | 14 | 4 | 14 | 32 | 44 |
|  | 3 | x |  | x |  | 15 | 36 | 11 | 17 | 5 | 17 | 32 | 49 |
|  | 4 | x | x |  |  | 13 | 26 | 14 | 16 | 4 | 13 | 32 | 44 |
|  | 5 | x |  |  |  | 12 | 32 | 14 | 21 | 4 | 16 | 31 | 50 |
|  | 6 |  |  |  |  | 13 | 21 | 32 | 69 | 0 | 9 | 161 | 344 |
| monkey E | 1 | x | x | x | x | 17 | 39 | 48 | 58 | 28 | 45 | 8 | 82 |
|  | 2 | x | x | x |  | 23 | 35 | 44 | 71 | 33 | 39 | 4 | 88 |
|  | 3 | x |  | x |  | 23 | 35 | 47 | 57 | 32 | 44 | 2 | 85 |
|  | 4 | x | x |  |  | 17 | 29 | 47 | 76 | 29 | 34 | 6 | 84 |
|  | 5 | x |  |  |  | 17 | 30 | 52 | 60 | 28 | 40 | 4 | 80 |
|  | 6 |  |  |  |  | 17 | 29 | 69 | 76 | 25 | 35 | 5 | 85 |

